# Text Mining of Disease-lifestyle Associations to Explain Comorbidities in Electronic Health Registries

**DOI:** 10.1101/168211

**Authors:** Lars Juhl Jensen

## Abstract

Mining of electronic health registries can reveal vast numbers of disease correlations (from hereon referred to as comorbidities for simplicity). However, the underlying causes can be hard to identify, in part because health registries usually do not record important lifestyle factors such as diet, substance consumption, and physical activity. To address this challenge, I developed a text-mining approach that uses dictionaries of diseases and lifestyle factors for named entity recognition and subsequently for co-occurrence extraction of disease–lifestyle associations from Medline. I show that this approach is able to extract many correct associations and provide proof-of-concept that these can provide plausible explanations for comorbidities observed in Swedish and Danish health registry data.

## 1. Introduction

In recent years, several groups have published systematic studies of simple comorbidities based on data from health registry data^1–2^ or text mining of the unstructured parts of electronic health records.^3–4^ More recent work has focused on taking into account the temporal ordering of the comorbid diseases and on combining pairwise comorbidities into longer disease trajectories.^5–6^

Regardless of the source of data, lifestyle is a major source of confounding factors in analyses of comorbidities. It is well known that diet, substance consumption and physical activity all influence the risk of getting many different diseases, and correlation is to be expected between any two diseases that are subject to the same confounding factors. As lifestyle is usually not recorded for patients who are not explicitly part of a study cohort, it is not possible to explicitly correct for this in the statistical analysis. Many of the identified comorbidities identified in large-scale studies of healthcare data may thus be trivially explained the patients’ lifestyles rather than by genetics.

It would be desirable to have a method to automatically identify the comorbidities most likely to be due to lifestyle before looking for a genetic explanation. This requires the construction of a knowledgebase of disease–lifestyle relationships grounded in a vocabulary or ontology of lifestyle factors. There are many efforts to create ontologies that capture a variety of aspects of lifestyle, such as environments (Environment Ontology^7^), exposure to risk factors (Risk Ontology^8^ and Exposome Explorer^9^), smoking (Cigarette Smoke Exposure Ontology^10^), and food (FoodOn, http://foodontology.github.io/foodon/). However, even if one were to unify all of these, important disease-associated factors like physical activity and socioeconomic status would not be captured.

Here, I provide initial proof-of-concept for text mining of disease–lifestyle associations from the biomedical literature and use of such associations to explain comorbidities in electronic health registry data. To this end, I have developed a draft dictionary of lifestyle factors and used it in conjunction with an existing disease dictionary to extract disease–lifestyle associations based on co-occurrences in Medline abstracts. By combining these associations with comorbidities found in electronic health registries, I show that this approach can identify plausible lifestyle-associated explanations for comorbidities. This suggests that text-mined disease–lifestyle associations can be used as a filter to help uncover comorbidities that are likely to have genetic causes.

## 2. Material and Methods

### 2.1. Compilation of a lifestyle dictionary

To identify candidate words for a lifestyle dictionary, I queried PubMed for “lifestyle[tiab] OR life-style[tiab]” and downloaded the PMIDs of the 75,761 matching abstracts (in the following referred to as lifestyle abstracts). Using a local copy of Medline, I next performed case-insensitive counting of the number of occurrences of all words within the lifestyle abstracts and within all abstracts. All words matching more than one million abstracts were removed to eliminate common English words. The remaining words were ranked using a simplified version of the scoring scheme previously used in the STRING^11^ and DISEASES^12^ databases. Whereas the full version of the scoring scheme calculates a weighted count of co-occurrences at the sentence and abstract levels, the simplified version counts only at the abstract level.

From the ranked list of candidate words, I manually compiled a lifestyle dictionary by first removing words such as lifestyle diseases and drugs, then grouping related words in a concept hierarchy, and finally adding additional word variants and synonyms. Whereas the resulting dictionary is by no means comprehensive, this approach should ensure that it includes most commonly used lifestyle descriptors and thereby that it is sufficient to provide proof-of-concept for disease–lifestyle association mining. This draft lifestyle dictionary is available for download at https://doi.org/10.6084/m9.figshare.5212603.

### 2.2. Named entity recognition of lifestyles and diseases

Next, I matched the dictionary of lifestyle terms against all Medline abstracts using an existing dictionary-based NER system, which is described in detail elsewhere^13^ and is available under BSD license at https://bitbucket.org/larsjuhljensen/tagger/. Briefly, the software is implemented in C++ and uses custom hashing and string-compare functions to perform flexible matching of dictionaries against text in a highly efficient manner. Good precision is ensured through the use of a manually curated, case-sensitive blacklist of unfortunate names, which would otherwise cause many false positives. To also identify disease names, I used the disease dictionary from the DISEASES^11^ databases, which was based on Disease Ontology^14^ and extended with mappings to the International Classification of Diseases (ICD) version 10^15^.

### 2.3. Associating diseases with lifestyles

To move from individual mentions of diseases and lifestyles to disease–lifestyle associations, I used a co-mention scoring scheme also used in the STRING^11^ and DISEASES^12^ databases. For completeness, the scoring scheme is briefly reiterated below.

The first step is to calculate a co-mention count of each disease with each lifestyle factor, using Eq. (1) to put more weight on co-mentions within sentences than across different sentences from the same abstract.

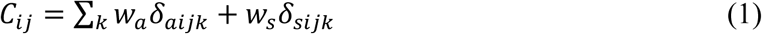

Here subscript *i* refers to a disease, subscript *j* to a lifestyle factor, and subscript *k* to an abstract. *C*_*ij*_ is the weighted co-mention count, *w*_*a*_ = 3 is the abstract weight, and *w*_*s*_ = 0.*2* is the sentence weight. *δ*_*ajik*_ and *δ*_*sjik*_ are 1 if *i* and *j* are co-mentioned in *k* within the same abstract and sentence, respectively, and 0 otherwise. Note that two entities that co-occur within a sentence will receive a total weight of *w*_*a*_ + *w*_*s*_.

The next step in the scoring is to compare the observed weighted co-mention count to what would be expected at random using Eq. (2).

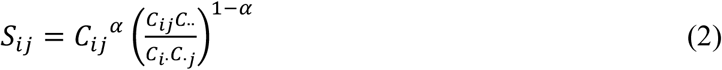

In this equation a subscript dots is shorthand for summing over all diseases or lifestyles. The parameter *α* = 0.6 specifies the relative weight put on the raw observed count vs. the observed/expected ratio. In this study all parameters were left at the optimal values, which were identified when developing STRING^11^ and found to perform well in DISEASES^12^ too.

The last step in the co-mention scoring is to normalize the *S*_*ij*_ scores to z-scores. This is done based on the global score distribution as previously published and filtered to retain only pairs with a z-score greater than 3.^12^ Finally, I filtered the set of associations to include only disease terms from ICD-10 level 3, because this is the level at which comorbidities were analyzed. The resulting sets of 1702 associations between ICD-10 codes and lifestyle factors are available for download at https://doi.org/10.6084/m9.figshare.5224501.

### 2.4. Retrieval of comorbidities from electronic health records

To explore if the text-mined disease–lifestyle associations can explain comorbidities, I compared them to two published sets of comorbidities between ICD-10 codes from Swedish and Danish health registry data, respectively. Comorbidities from the HuDiNe study^2^ were not used because they are coded in ICD-9-CM, which cannot be easily mapped to ICD-10.

The first set consists of comorbidities identified by Dalianis and coworkers in the Stockholm Electronic Patient Record (SEPR) corpus.^1^ To avoid sex as a confounding factor, the female- and male-specific comorbidity data were retrieved from https://www2.dsv.su.se/comorbidityview-demo/. However, the age distributions of ICD-10 codes were not corrected for, as sufficiently finely resolved data were not available. I calculated the relative risk of each comorbidity from the raw count data for each sex, and kept only pairs of level-3 ICD-10 codes which for at least one sex showed a relative risk greater than 10 and were supported by more than 20 patients. This resulted in a set of 902 comorbidities in total.

The second set consists of directional comorbidities identified by Jensen and coworkers in the Danish National Patient Registry (NPR).^4^ In contrast to the SEPR dataset, both the sex and age distributions of the codes were explicitly corrected for in the published statistical analysis and combined results, including relative risk, were reported. Other important differences are that the NPR comorbidities were tested for statistical significance, both for the correlation between codes and for their temporal directionality. Filtering the dataset to consider only pairs with relative risk greater than 10 supported by at least 20 patients yielded 328 directional comorbidities between level-3 ICD-10 codes.

### 2.5. Visualization of the disease–lifestyle network

To visualize the complex interplay of diseases and lifestyle factors I imported both comorbidity networks and the disease–lifestyle associations into Cytoscape. I next extracted a module around each life style factor in the network, including only diseases that were comorbid with at least one other disease associated with the same lifestyle factor according to the SEPR or NPR data.

## 3. Results and Discussion

### 3.1. Assessment of extracted disease–lifestyle associations

An important step after performing text mining of disease–lifestyle associations is to assess if the approach succeeds in extracting known associations. However, in the absence of a manually curated resource of such associations, this is difficult to do in a formal manner. Instead, I opted to inspect the top diseases associated with each of ten lifestyle factors. These include consumption of substances (alcohol, caffeine, recreational drugs, and tobacco), eating habits (binge eating, dieting, and snacking), macronutrients (carbohydrates and fats), and physical activity. The top-10 diseases (i.e. ICD-10 codes) for each of these lifestyle factors are shown in Table 1.

**Table 1.** Top-10 diseases associated with each of ten lifestyle factors.

| Lifestyle factor | Diseases (ICD-10 level 3) |
| --- | --- |
| Alcohol consumption | K70, K76, E75, K74, E88, E90, D48, C80, E74, E14 |
| Caffeine consumption | D48, C80, G96, E88, E90, E74 |
| Recreational drug use | F20, F90, G96, F41, F60 |
| Tobacco consumption | D48, C80, E14, E74, E88, E90, J85, J98, J44, J45 |
| Binge eating | F50 |
| Dieting | F50 |
| Snacking | K02, K03 |
| Carbohydrate | E88, E90, E74, E14, D48, C80, E16, E35, E11, K02 |
| Fat | E88, E90, E74, E14, D48, I70, C80, E78, E35, E11 |
| Physical activity | E88, E90, E74, E14, G96, M13, I67, D48, C80, I51 |

The first thing that stands out is a set of five diseases that are all related to consumption of alcohol, caffeine, recreational drugs, tobacco, carbohydrate, and fat as well as to physical activity. The codes fall into two groups, namely unspecified cancers/neoplasms and broad metabolic disorders: malignant neoplasm, without specification of site (C80), neoplasm of uncertain or unknown behavior of other and unspecified sites (D48), other disorders of carbohydrate metabolism (E74), other metabolic disorders (E88), and nutritional and metabolic disorders in diseases classified elsewhere (E90). While not very informative, these codes are consistent with the extensive literature on the impact of lifestyle on the risk of developing cancer and metabolic disorders. However, these will not be discussed further below due to their unspecific nature.

Binge eating and dieting are both exclusively associated with eating disorders (F50). Only two diseases were found to be associated with snacking, namely dental caries (K02) and other diseases of hard tissues of teeth (K03), both of which have obvious associations with eating candy. Dental caries also associates with carbohydrate intake, which is furthermore linked to diabetes mellitus (E11 and E14) and other diseases related to the pancreas (E16) and other endocrine glands (E35). Fat intake and (lack of) physical exercise are linked to many of the same diseases as well as to atherosclerosis (I70) and vascular diseases (I51 and I63), respectively. The link between physical activity and disorders of central nervous system (G96) can be viewed as a false positive, as the co-mentioning is primarily due to cerebrospinal diseases that prevent physical activity.

In addition to the five unspecific ICD-10 codes mentioned above, including cancer, tobacco consumption is specifically associated with pulmonary disorders (J44, J45, J85, and J98). Recreational drug use was found to associate with a wide range of mental and behavioral disorders (F20, F41, F60, and F90). Again, disorders of central nervous system (G96) showed up as what can be viewed as a false positive, the reason being medical uses of cannabinoids. Finally, caffeine is co-mentioned with cancers, metabolic disorders, and central nervous system disorders (G96); however, inspection of the underlying abstracts revealed that most studies found caffeine to have no influence on disease risk.

This inspection of the diseases most strongly associated with ten lifestyle factors suggests that the simple text-mining approach presented here is able to extract many correct disease–lifestyle associations from literature with a modest error rate.

### 3.2. Explaining comorbidities through shared lifestyle factors

The next step is to combine the text-mined disease–lifestyle associations with comorbidity data to see if the former can explain the latter. A small network exemplifying this is shown in Fig. 1.

**Fig. 1.**
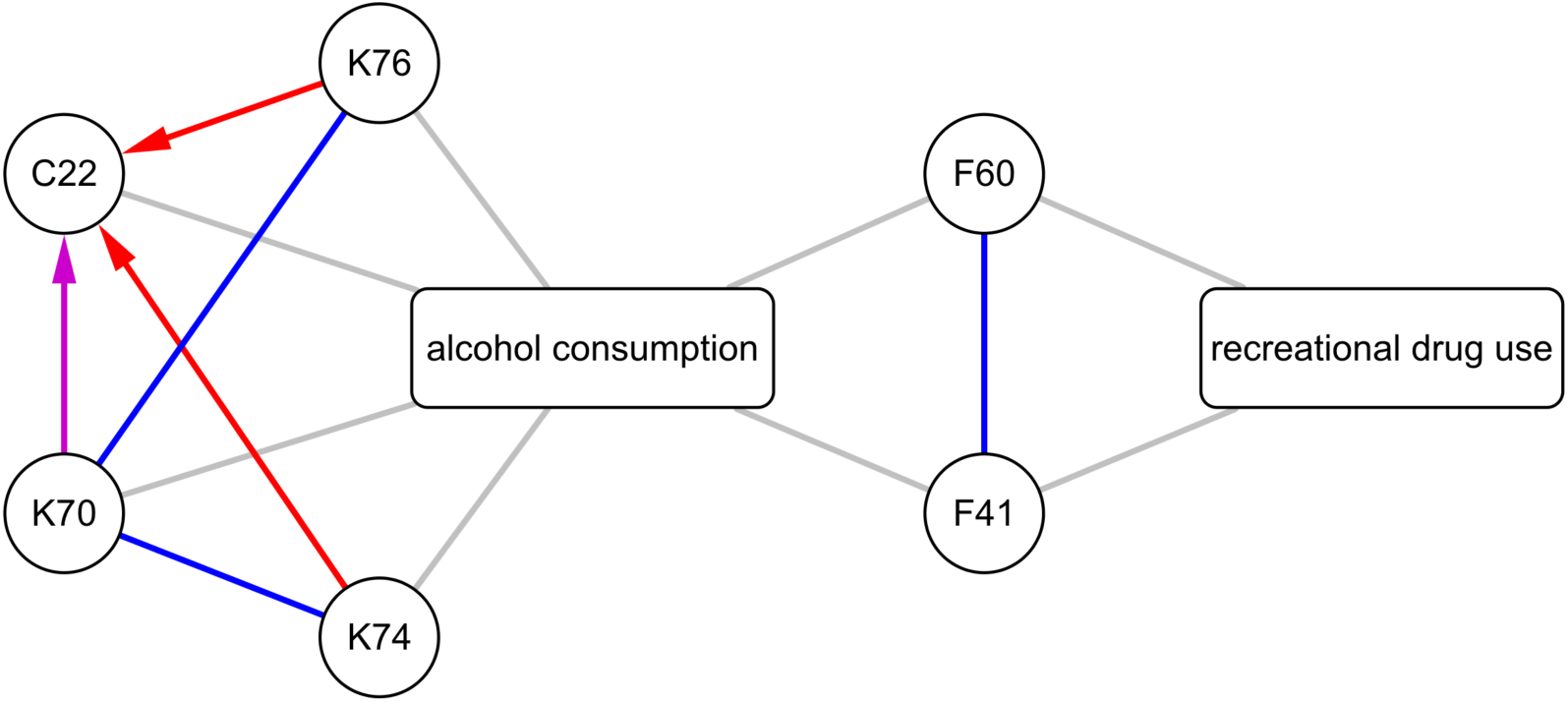
The network shows the text-mined associations (gray lines) between two lifestyle factors and six diseases, namely alcoholic liver disease (K70), liver fibrosis/cirrhosis (K74), other liver diseases (K76), liver cancer (C22), anxiety disorders (F41), and personality disorders (F60). The blue lines are undirected comorbidities from the Swedish SEPR data, the red arrows are directed comorbidities from the Danish NPR data, and the purple arrow is a comorbidity found in both.

To quantify if the lifestyle factors can indeed explain comorbidities, I counted the number of comorbidities that could be explained by one or more lifestyle terms, not considering the four broadest terms (lifestyle, physical fitness, sociodemographic factor, and substance consumption). Of the 902 undirected comorbidities from the SEPR data, 46 could be explained by one or more lifestyle factors that are strongly associated with both diseases, which is exactly twice as many as in a randomized network (P < 1%, Fisher’s exact test). This shows that comorbid diseases are indeed tend to be associated with common lifestyle factors, although the proof-of-concept approach presented here only finds such explanations for a small percentage of disease pairs.

Performing the same analysis on the 328 temporally directed comorbidities from NPR data explained 32 disease pairs in the real network and 26 in the randomized network (P > 10%, Fisher’s exact test). This suggests the requirement of temporal directionality eliminates many of the comorbidities between diseases with common lifestyle factors.

### 3.3. Future work

Since the work presented here is only intended to show the feasibility and utility of text mining disease–lifestyle associations, there is obviously room for improvement on all fronts. Even then, the results at hand show sufficient promise to warrant taking on the many open challenges.

To capture lifestyle factors in a structured manner, it would clearly be preferable ground the dictionary in one or more ontologies; however, doing so will be a very time-consuming endeavor compared to the *ad hoc* approach used here. Once accomplished, the NER approach needs to be benchmarked on a manually annotated text corpus, the construction of will again involve a nontrivial amount of work. The latter will also allow training of machine-learning-based NER methods and comparison of their performance to the dictionary-based approach used here.

Many of the same considerations apply to the subsequent relation extraction task. Without a manually curated corpus or database of disease–lifestyle associations, it is difficult to quantify how well a method works or say if a different approach would be better. As noted earlier, the disease–lifestyle associations extracted here are a mix of factors that increase and decrease the risk of getting certain diseases as well as factors that have been extensively studied but found to have no impact (e.g. caffeine). Ideally, one would want to also benchmark how well the indirect disease–lifestyle–disease associations explain the observed comorbidities, which will likely prove even more challenging. Finally, although I expect one can enrich for comorbidities with a genetic cause by filtering out those that are likely explained by lifestyle, this remains to be demonstrated.

## 4. Conclusions

Data mining of electronic health registries shows great promise for improving our understanding of comorbidities and disease progression. However, it remains a major challenge to account for the numerous confounding factors, many of which relate to the lifestyles of patients. As the registries do generally not record such patient-level information, this cannot be corrected for in the statistical analyses. The result is that comorbidities with genetic explanations become buried among vast numbers of trivial comorbidities that are explainable by lifestyle. Being able to separate the two would be an important step towards being able to act upon discovered comorbidities in a clinical setting.

The lifestyle factors implicated in many individual diseases are well known and described in the biomedical literature, but have not been systematically recorded in a structured, searchable database. Here, I present a proof-of-concept that such a resource can be constructed through text mining of biomedical abstracts. Through comparison with published analyses of Swedish and Danish medical registries, I also show that disease–lifestyle associations can provide plausible explanations for comorbidities observed in Swedish and Danish health registry data. The approach described in this work may help improve our understanding of disease trajectories and explain lifestyle causes for comorbidities in individual patients.

## Acknowledgments

Thanks to David Westergaard for providing the Disease Ontology to ICD-10 mapping. This work was supported by the Novo Nordisk Foundation (grant agreement NNF14CC0001) and the European Commission under the European Union’s Horizon 2020 research and innovation programme (grant agreement 668031).

